# Distribution of disease-causing germline mutations in coiled-coils suggests essential role of their N-terminal region

**DOI:** 10.1101/2020.04.07.029165

**Authors:** Zsofia E. Kalman, Bálint Mészáros, Zoltán Gáspári, Laszlo Dobson

## Abstract

Next-generation sequencing resulted in the identification of a huge number of naturally occurring variations in human proteins. The correct interpretation of the functional effects of these variations necessitates the understanding of how they modulate protein structure. Coiled-coils are α-helical structures responsible for a diverse range of functions, but most importantly, they facilitate the structural organization of macromolecular scaffolds *via* oligomerization. In this study, we analyzed a comprehensive set of disease-associated germline mutations in coiled-coil structures. Our results highlight the essential role of residues near the N-terminal part of coiled-coil regions, possibly critical for superhelix assembly and folding in some cases. We also show that coiled-coils of different oligomerization states exhibit characteristically distinct patterns of disease-causing mutations. Our study provides structural and functional explanations on how disease emerges through the mutation of these structural motifs.

## Introduction

Advances in sequencing resulted in the identification of a huge number of different Single Nucleotide Variations (SNVs) of various genomic positions among individuals in healthy and disease states. Variations can impact proteins on several levels, ranging from polymorphisms (PMs) with negligible effect on fitness to lethal mutations through increasingly strong phenotypes. According to their origin, disease-causing genetic alterations can be broadly categorized into either somatic or germline mutations. Somatic mutations, most notably responsible for tumorigenesis, are confined to the cell they originated in and its daughter cells.

In accord, their phenotypic change can be extreme with abolishing cell-cycle control, escaping apoptosis and achieving cellular immortality. In contrast, disease-associated germline mutations (DMs) persist in the whole organism and are transmitted from generation to generation. Thus, DMs cause relatively weak changes in phenotypes, yet they still have a noticeable negative impact on the quality of life.

Coiled-coils are oligomeric helical structural units in proteins connected to a wide range of functions. Several coiled coil proteins have been shown to have catalytic activity^1^ or undergo oligomerization, yet their most common functions arise directly from their structure: they are molecular spacers separating or connecting domains^2^. They can bridge large distances and connect proteins of different sides of large structural organizations like the postsynaptic density where the Homer coiled coil serves as a direct connection between the plasma membrane and intracellular proteins through EVH1 domains binding to various scaffolding partners^3^.

Coiled-coils are α-helical domains consisting of two or more helices packed together in a specific knobs-into-holes manner^4^, with interhelical interactions playing a dominant role in folding^5^. Coiled-coils can have parallel or antiparallel arrangement, and they can be formed by intrachain interaction of the same subunit, or by interchain bonds between distinct polypeptide chains^6^. Regardless of their oligomerization state, the main forces driving their interactions are the formation of hydrophobic contacts at their inside, often supported by electrostatic interactions aiding the stability from the outside^7^. Coiled-coil residues can be classified into register positions according to their role in complex formation with the opposing helices: the most common repeat pattern is heptad (‘abcdefg’), with ‘a’ and ‘d’ positions being responsible for hydrophobic interactions, while ‘e’ and ‘g’ positions contain (oppositely) charged residues^8^. Notably, other variants (e.g., hendecads are 11 residue repeats) were also discovered^9^. Folding studies of selected coiled-coils indicated the importance of a specific segment, the trigger sequence, that is required for initiating the proper interaction between the helices^10^. However, it is not yet entirely clear whether specific sequence patterns are required for assembly, or the accumulation of interaction promoting residues at critical positions generally aids coiled-coil formation.

In recent decades many studies addressed how DMs perturb protein structure. The majority of significantly occurring structural elements (e.g., transmembrane ^11^ and intrinsically disordered protein regions^12^), as well as various structurally distinct functional regions (e.g., protein-protein interfaces, buried domains^13^) were analyzed in detail. However, coiled-coils are a largely understudied class in this respect with only individual cases discussed. To our knowledge, only one large-scale study has been published, highlighting the critical role of register positions and pointing out mutations frequently associated with pleiotropy^14^. In this study we integrate multiple prediction algorithms and structural information for an in-depth analysis to assess how non-synonymous disease-associated germline mutations affect coiled-coil structures and thereby their functions. Our work revealed that disrupting hydrophobic and electrostatic interactions impairs coiled-coil structure and disease-associated mutations accumulate near the N-terminal of coiled-coil regions. We also showed that even if their destabilizing effect is small, DMs are enriched in antiparallel homodimer coiled-coils. On one hand, understanding how these variations modulate the structure and function of proteins may improve prediction algorithms. On the other hand, the rational coiled-coil design can be achieved through a detailed understanding of the sequence-structure relationship^15^. However, a missing piece of the puzzle is how DMs perturb coiled-coil structures.

## Results

### DMs primarily target coiled-coil segment containing proteins outside the coiled-coil region

To obtain an overall picture of how disease-associated mutations (DM) and coiled-coils are related, we determined the number of proteins in the human proteome that contain coiled-coil regions and at least one DM or polymorphism (PM). To define coiled-coil regions, we used four different predictors (Deepcoil, Marcoil, Ncoils and Paircoil). We calculated all statistics separately for each method (detailed in the supplementary material) and also for the consensus of the predictions.

In general, PMs have a higher overall abundance in the human proteome than DMs, as observed in many previous studies. At the protein level, there is a slight increase in the frequency of DMs in coiled-coil containing proteins compared to other proteins in the proteome, yielding an odds ratio of 1.03 against PMs (Supplementary Figure 1). Notably, prediction methods exhibit some inconsistency regarding these statistics: DeepCoil predicts less coiled-coils and produces a negative odds ratio, while the other three methods show the inverse trend. Surprisingly in the light of the above observation, coiled-coil regions are depleted in DMs compared to non-coiled-coil residues, agreed by all prediction methods. A possible explanation for this phenomenon might be that coiled-coils are generally less vulnerable to harmful variations. However, these proteins carry essential functions as they are more frequently occupied by DMs.

### Coiled-coils are often perturbed by DMs affecting charged residues

The main driving force of protein domain folding and stability is achieved through hydrophobic interactions. Coiled-coils are special structural units where the balanced contribution of hydrophobic interactions and electrostatic interactions aid the stability together. This is reflected by the different amino acid preferences of the different positions in the heptad repeat unit, corresponding to the distinct spatial position and role of these within the superhelical structure. To assess how residues are affected by variations, we grouped amino acids based on their basic physico-chemical features (positive: HKR, negative: DE, hydrophobic: AILMV, other: CFGNPQSTWY), then we calculated the log ratio of the substitutions observed in DMs.

Figure 1 shows preferred residue type changes in coiled-coils relative to other proteins of the proteome. We calculated the relative frequency of amino acid substitutions in the coiled-coil regions, and in the proteome, then calculated the log ratio of substitution frequencies. According to our results hydrophobic residues are targeted in similar proportions. However, in the case of coiled-coils, charged residues aiding electrostatic interactions are much more frequently affected by DMs (Figure 1/right).

**Figure 1:**
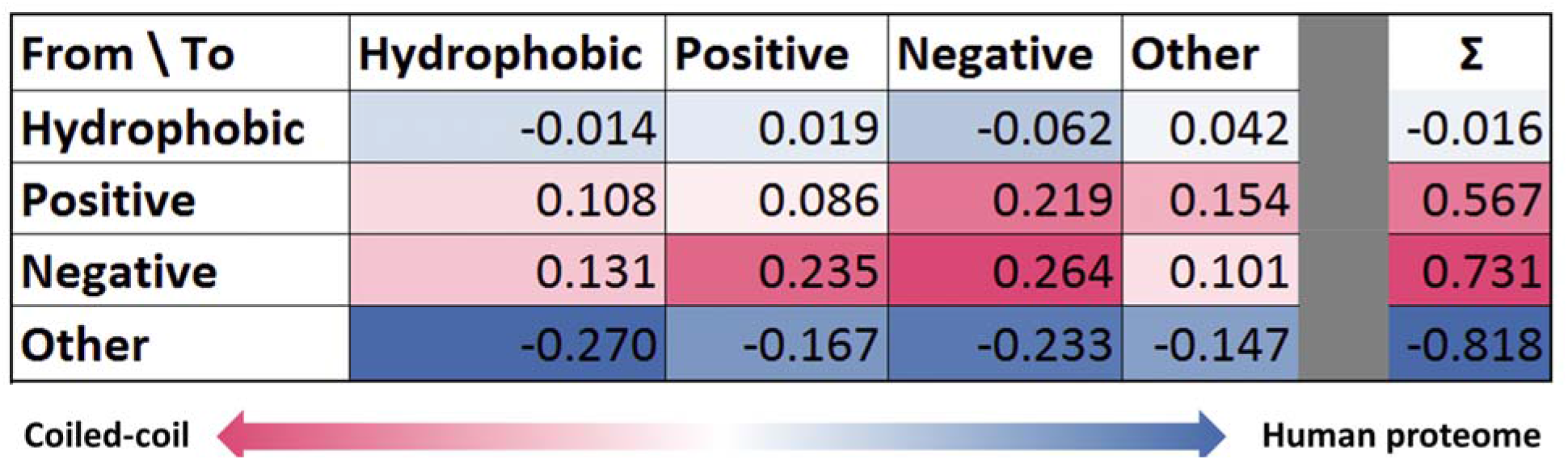
Amino acid changes in coiled-coils. Left) Residue change preferences by DMs in the proteome (negative values, also marked with the shades of blue) and in coiled-coil regions (positive values, also marked with the shades of red). Values show the logarithm of ratio of DMs changing given residues types Right) Targeted residue type preferences by DMs in the proteome (negative values) and in coiled-coil regions (positive values).

In contrast, several residue types indispensable for stable domain structure (e.g., cysteines forming disulfide bridges) do not influence coiled-coil formation, thus their replacement does not cause stability problems (Figure 1/left, for more details see Supplementary Figure 2). The most prevalent changes in coiled-coil regions by DMs are replacements by oppositely charged residues. Interestingly, the negatively charged Glu and Asp are generally not interchangeable residues in coiled-coils, in contrast to the positively charged residues Lys and Arg. In coiled-coils DMs most likely target A,E,I,K,L,M,N and Q residues, as opposed to C, G and P residues being more often targeted in other proteins in the proteome (Supplementary Figure 2).

### DMs accumulate at the N-terminal region of coiled-coils

We investigated the distribution of variations in coiled-coils, considering their coverage, abundance in the N-terminal region, and coiled-coil length. We divided coiled-coil regions into five equal parts, and calculated the proportion of variations in these parts. Although the first half of the sequences contain slightly more DMs, the difference is not significant compared to PMs (Supplementary Figure 3).

To reduce bias originating from the varied length of coiled-coils, we performed the enrichment calculation considering only the first 28 residues of all coiled-coils, this time by dividing sequences into four equal parts, i.e., using seven residue bins - keeping in mind that predictors were optimized for heptad repeats (Figure 2/A). Using this approach, the accumulation of DMs at the N-terminal became visible, showing a monotonous decline of DMs towards the C-terminal.

**Figure 2:**
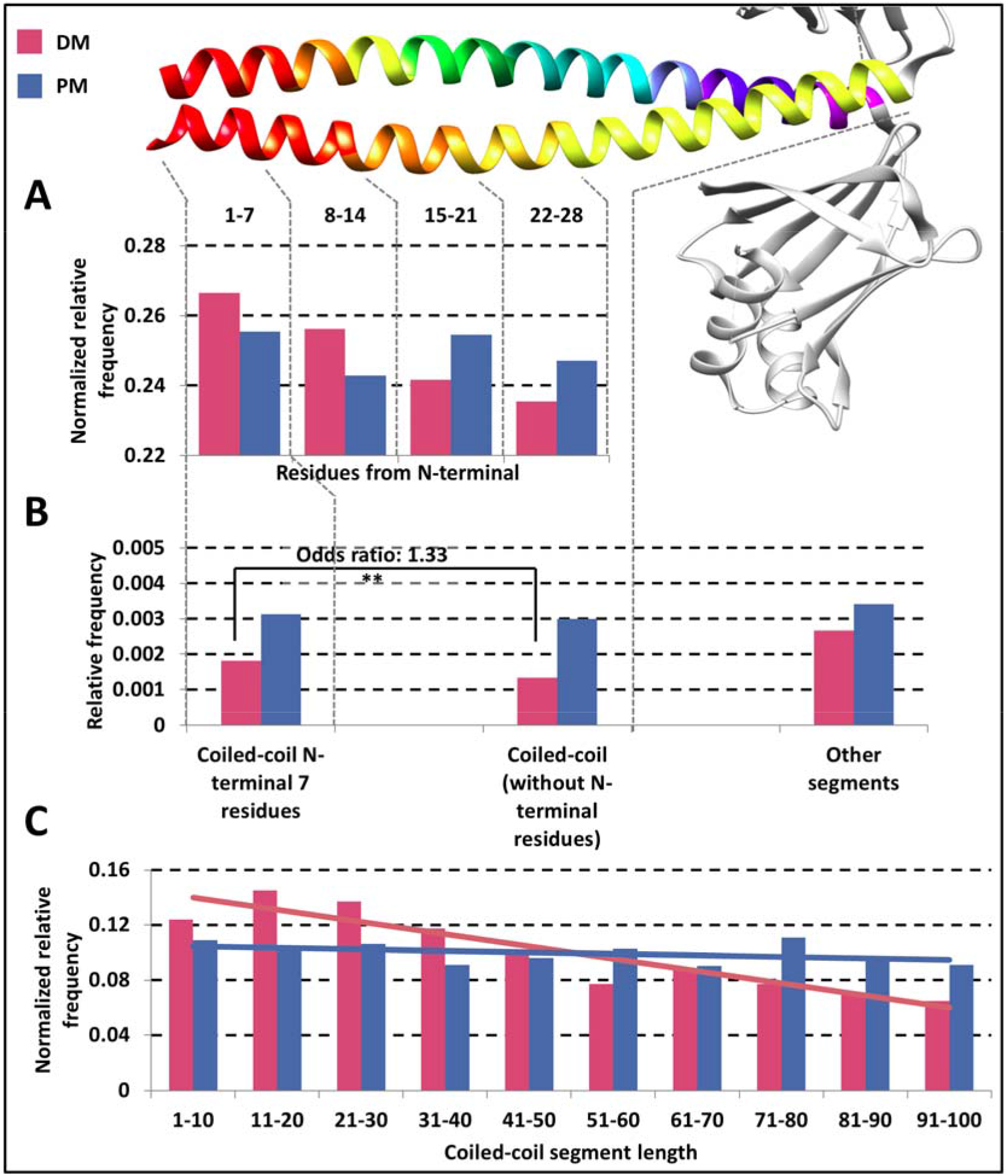
Variations along coiled-coil segments. Distribution of variations in the sequence. A) X-axis shows the coverage of the N-terminal of coiled-coils. B) Relative frequency of coiled-coil residues targeted at N-terminal of the coiled-coil, other coiled-coil residues and other segments of proteins, respectively. C) Distributions of variations in coiled-coils with different lengths (linear trend lines were aligned to the data). Red: DMs; blue: PMs.

To demonstrate that the first seven residues of coiled-coils contain significantly more DMs compared to the rest of the coiled-coil regions, we counted the number of DMs and PMs in the first seven residues of coiled-coils and in their succeeding part. The result is significant (χ^2^ test, p<0.01), and the odds ratio between DMs and PMs is 1.33 (Figure 2/B).

Notably, this effect is strong enough to influence the distribution of variations considering coiled-coils with different lengths. The relative frequency of DMs is significantly higher in shorter coiled-coils (Figure 2/C), as they utilize most of their residues as “N-terminal” segment that may contribute to stability, while in longer coiled-coils, other residues have a lesser role in sustaining the complex form. In contrast, PMs show uniform distribution in coiled-coil regions with different lengths. Notably, it is arguable whether very short predicted coiled-coil segments (below 10 residues) are biologically relevant, however we did not want to tailor prediction outputs. Moreover, omitting the first bin only further strengthens our result.

We performed the same analysis around C-terminal residues, however according to our results the accumulation of DMs is only detectable at the N-terminal region (data not shown).

### Oligomerization state affects which register positions are vulnerable

The periodic property of coiled-coils enables a position type classification of residues, grouping amino acid positions based on their location in the helix, uncovering preferred physico-chemical features and interaction types. Regardless of oligomerization state, residues at “a” and “d” positions are often hydrophobic and face each other, forming the core of the complex, while “e” and “g” residues may be charged and promote stability via electrostatic interactions on the outer face of the superhelical structure.

We analyzed the distribution of variations on the different positions, considering different oligomerization states. As expected from early results, PMs are more abundant in every heptad position. However, considering DMs only, residues falling into “stabilizing” positions are more vulnerable to variations (Figure 3, left). Interestingly, residue type changes affect the heptad positions differently: replacement of amino acids in “a” and “d” positions likely perturb the structure, even when the substitution seems conserving on the basis of physicochemical properties (i.e., variation replacing a hydrophobic residue to another one is also often harmful). In contrast, “e” and “g” positions seem to be slightly more resilient, and residue type change (i.e., charge change) is more often required to disrupt the structure. PMs change the residue type to a lesser extent.

**Figure 3:**
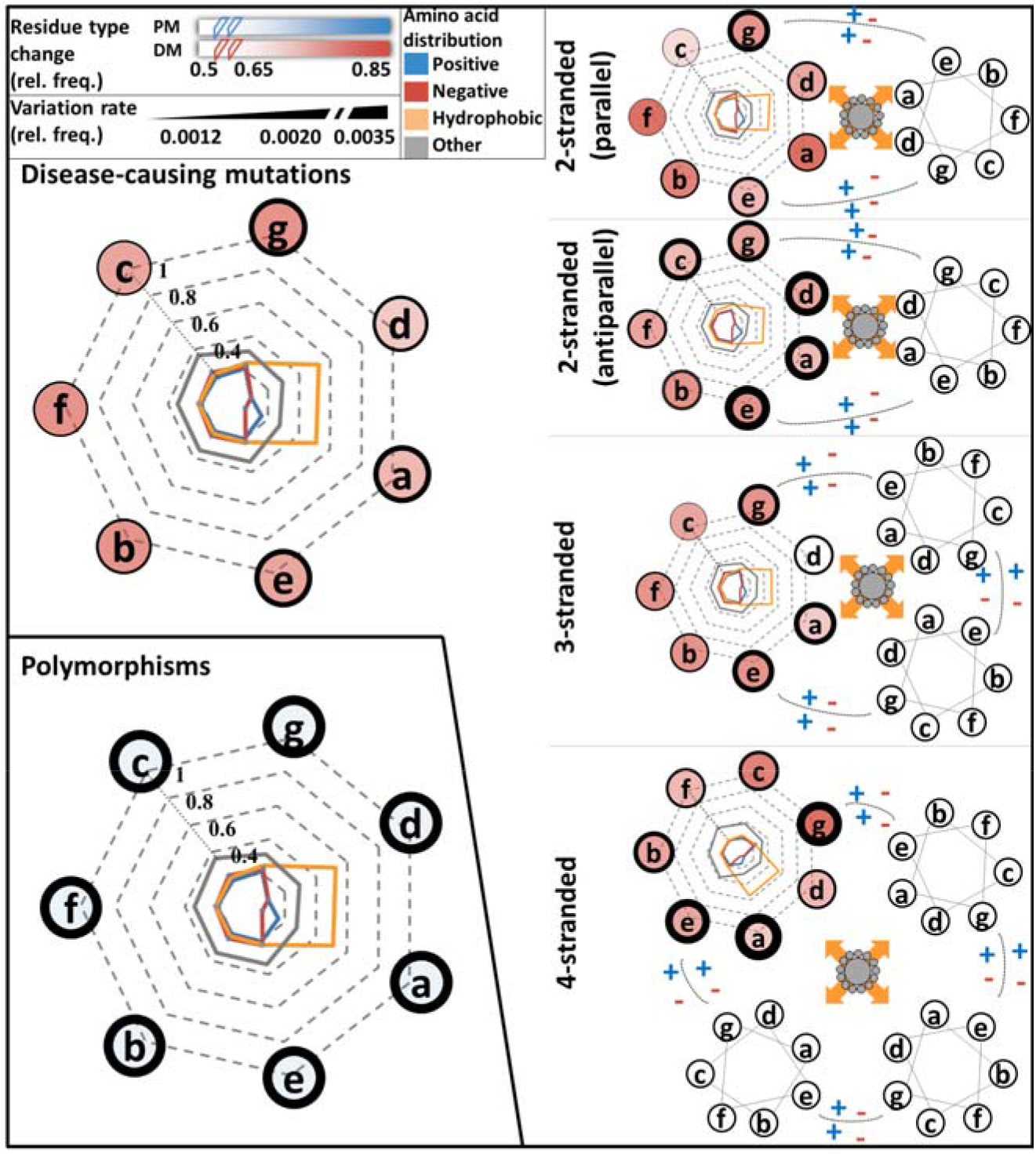
Distribution of variations in coiled-coils. Amino acids were grouped according to their physico-chemical properties (positive: HKR, negative: DE, small hydrophobic: AILMV, other: CFGNPQSTWY). Radars represent the amino acid distributions in different positions. Line thickness around positions is proportional to variation frequency. The opacity of positions is proportional to the rate of variations changing the physico-chemical features of the targeted residue. Left) DMs and PMs in coiled-coils. Middle) DMs in coiled-coils with different oligomerization states.

The oligomerization state of the coiled-coil also affects its vulnerability. In general, antiparallel formations (both dimers and tetramers) are slightly more preferred targets of DMs. Oligomerization also influences which positions are modulated (Figure 3, right): “e” and “g” (charged) positions are more often affected by DMs in parallel dimers, while “a” and “d” (hydrophobic) positions are primarily targeted in antiparallel dimers. Hydrophobic interaction promoting positions are less likely to be targeted by DMs in parallel dimers. When these positions are mutated, the mutation changes the type of the residue in almost every case, showing an opposite trend compared to other oligomerization modes.

Variations in trimeric and tetrameric coiled-coils are similar: in these cases, structures are most often perturbed via amino acids in “a”, “g” and “e” positions and also often replace residue type. DMs on “d” positions are rare.

In general, there seems to be an opposing trend, that in positions where the DM frequency is lower, any change can carry disease, while in positions where the DM frequency is higher, the mutations more likely change the physico-chemical property of the residue.

### Structure analysis reveals most DMs occur in homooligomeric coiled-coils with a subtle destabilizing effect

To gain detailed insights on how DMs perturb the formation of coiled-coils, we searched for structures in the PDB and identified coiled-coil segments using SOCKET. Although the number of variations falling into characterized coiled-coil structures is rather low, and sometimes insufficient for performing reliable statistical tests to draw convincing conclusions, such analysis can open prospects to recognize interesting trends.

First, we analyzed the distribution of DMs in different heptad positions. As the number of cases was low, we classified the positions into three categories: responsible for hydrophobic stabilization (a,d), electrostatic stabilization (e,g) and outward facing/solvent exposed (b,c,f). Disease-associated mutations are enriched on residues responsible for forming the hydrophobic core of coiled-coils, have nearly the same occurrence as PMs in positions reserved for charged residues, and show decline on outward facing residues (Figure 4/A). Although at first glance this does not seem to confirm prediction data where DMs have a higher frequency on ‘e’ and ‘g’ positions. However, this discrepancy is due to the very different composition of the two datasets with regard to oligomerization state: the most prevalent class of structures are two-stranded antiparallel coiled-coils, the only class where mutations on hydrophobic positions dominate in prediction data too (Figure 3).

**Figure 4:**
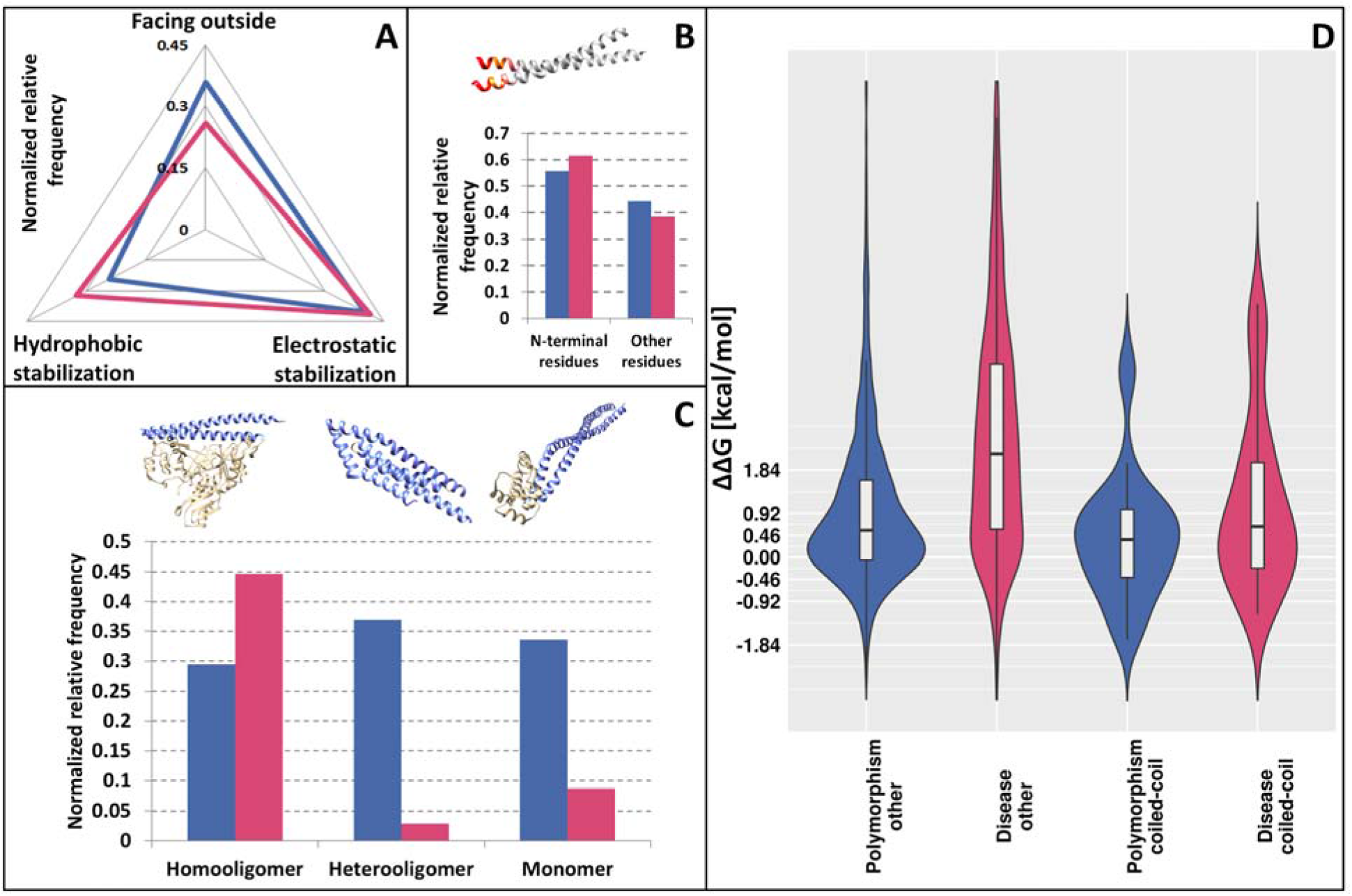
Distribution of variation on PDB structures. A) Distribution of variations based on the register position types. B) Distribution of variations in the N-terminal seven residues and in other segments of coiled-coils. C) Distribution of variations according to the oligomerization state of structures. D) Energy change distributions in coiled-coils (right) and in other proteins from the human proteome (left). Red: DMs; blue: PMs.

Next, we investigated whether N-terminal segments of the coiled-coils gather more variations. Although both types of variations (PMs and DMs) seem to accumulate in the first seven residues of coiled-coils, the two kinds of variations exhibit an opposing trend, with a higher frequency of DMs around the N-terminal and PMs in other residues (Figure 4/B). Moreover, while PM data is not significant, the distribution of DMs is slightly significant according to the χ^2^ test (p<0.1).

Sequence data alone can be rather difficult to utilize for defining the monomeric or oligomeric state of coiled-coil assemblies, and predictions are also limited in detecting how many strands the coiled-coils are composed of. However, from structural data we can readily classify coiled-coils as monomers (both strands are part of the same protein, typically antiparallel coiled coils with a short linker between the two helices), homooligomers (interaction of identical proteins) or heterooligomers (interaction between different proteins). While PMs show a uniform distribution among these classes, DMs mainly occur in homooligomers (Figure 4/C). The rationale behind this can be that a single mutation might (but in a heterozygous case, not necessarily) affect multiple constituent helices simultaneously, so their effect is instantly multiplied, in contrast to heterooligomers and monomers where the interacting partner/segment does not amplify the impact of the mutation.

The energy change calculated by the introduced mutation can be used as an approximation of the contribution of a mutation to the overall stability of the coiled coil. Figure 4/D shows the calculated energy changes upon mutation in the proteome and in coiled-coil structures. Generally, the mean energetic contribution of PMs can outline the range of changes a protein can tolerate without damage. In both cases (proteome, and coiled-coil proteins), DMs have an average higher ΔΔG. However, in the case of coiled-coils, despite keeping the same trend, both variation types seem to have a lower effect compared to those of other proteins of the proteome.

Further context can be added by the joint analysis of structural data. Figure 5 shows how DMs are distributed according to their features. Heptad positions with the highest standard deviation corresponding to energetic changes (positions ‘b’, ‘d’, ‘f’) exhibit the most heterogeneous distribution of different types of coiled-coils. Mutations in positions contributing to the hydrophobic core of coiled-coils (‘a’ and ‘d’), as well as the most outward facing residue type (‘f’) available for interactions operate with the most destabilizing energy changes: mutations here likely have more critical effect compared to other positions where there is more (spatial) room for substitutions. Mutations on ‘a’ position more likely affect two-stranded antiparallel coiled-coils (77%; 10 mutations affect two-stranded antiparallel coiled-coils out of total 13 mutations on position ‘a’), which is interestingly not true for the other hydrophobic residue promoting position ‘d’ (22%), also confirmed by the more comprehensive prediction data. Negative (stabilizing) energetic change is somewhat more likely occurring in homooligomeric coiled-coils (80%), with most cases on position ‘e’. We mapped the variations to only one chain of PDB structures, thus the real energetic contribution of a mutation may be even more stabilizing in homooligomers, abolishing transient interactions. In contrast, most mutations affecting monomeric coiled-coils are definitely highly destabilizing (94%), suggesting greater energetic effect is required to disrupt the overall structure that also includes intrachain interactions outside the coiled-coil, in contrast to mutations of complexes where coiled-coil interchain interactions are the only forces keeping the complex together. Last, but not least, destabilizing mutations more frequently occur in the possible N-terminal regions (57%), compared to stabilizing mutation (33%).

**Figure 5:**
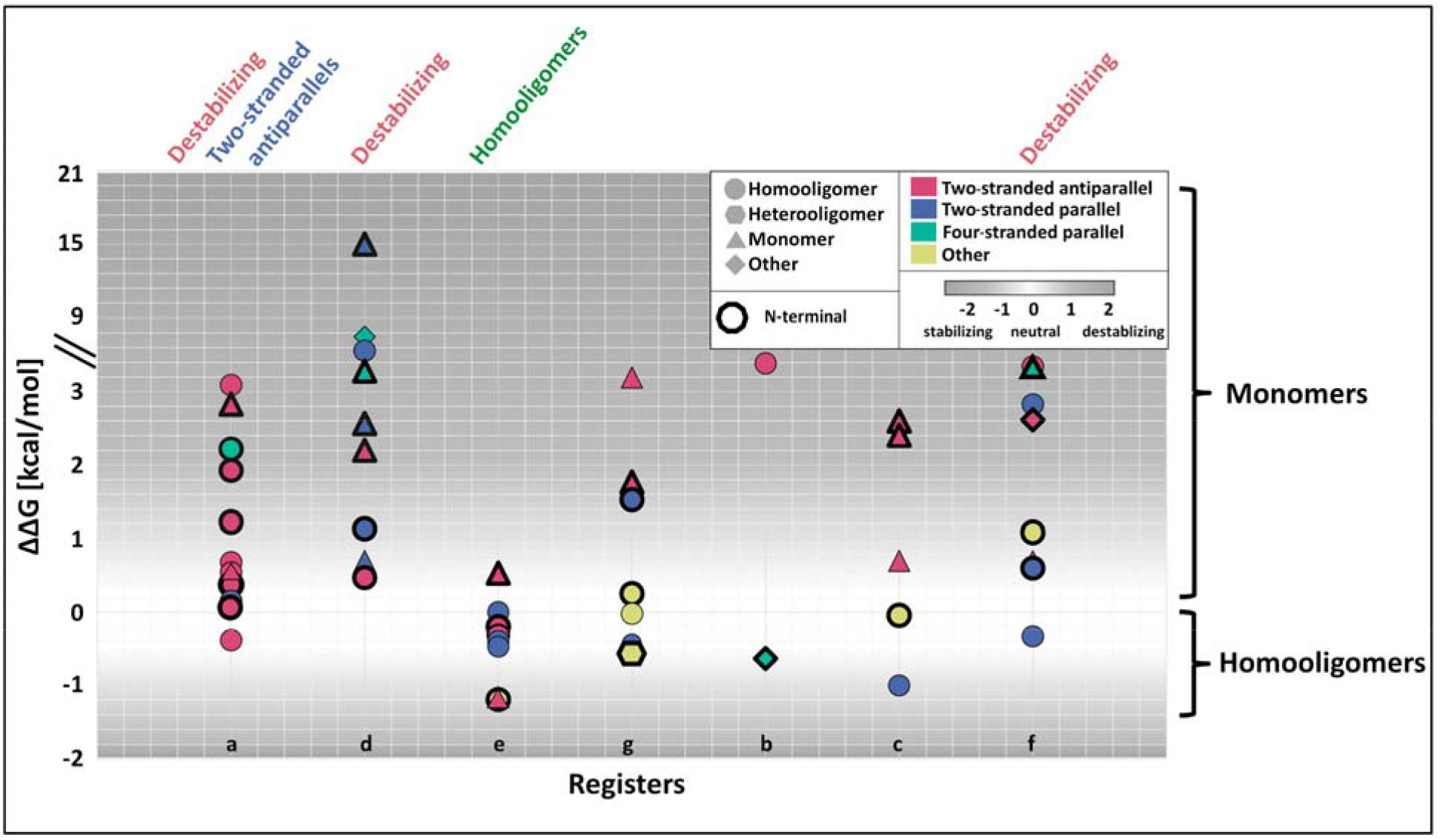
Distribution of DMs with respect to various structural features. Y-axis shows the calculated energy change caused by mutations. X-axis shows the register position DMs fall into. Colors and shapes highlight further properties (see legend).

### Impaired coiled-coils are mostly associated with central nervous system diseases

To get a better picture of what types of diseases emerge from the analyzed variations, we used DiseaseOntology to group diseases into categories based on their MIM identifier. Such ontologies often have some level of annotation bias, as the included proteins are not necessarily annotated to the same extent. To avoid drawing false conclusions we did not calculate p-values, and only counted the number of occurrences (and also highlighted the expected number of occurrences for comparison). Table 1 shows the most commonly occurring disease classes, where a DM directly disrupts the structure of a coiled-coil. According to our analysis, the most enriched disease terms are skin diseases, muscular diseases, carbohydrate metabolic diseases, and central nervous system diseases. Coiled-coil DMs are also enriched in more general terms, such as integumentary system diseases, musculoskeletal system diseases, and nervous system diseases. There is also a subtle abundance of conditions related to metabolic diseases that can be acquired according to the annotation.

**Table 1:**
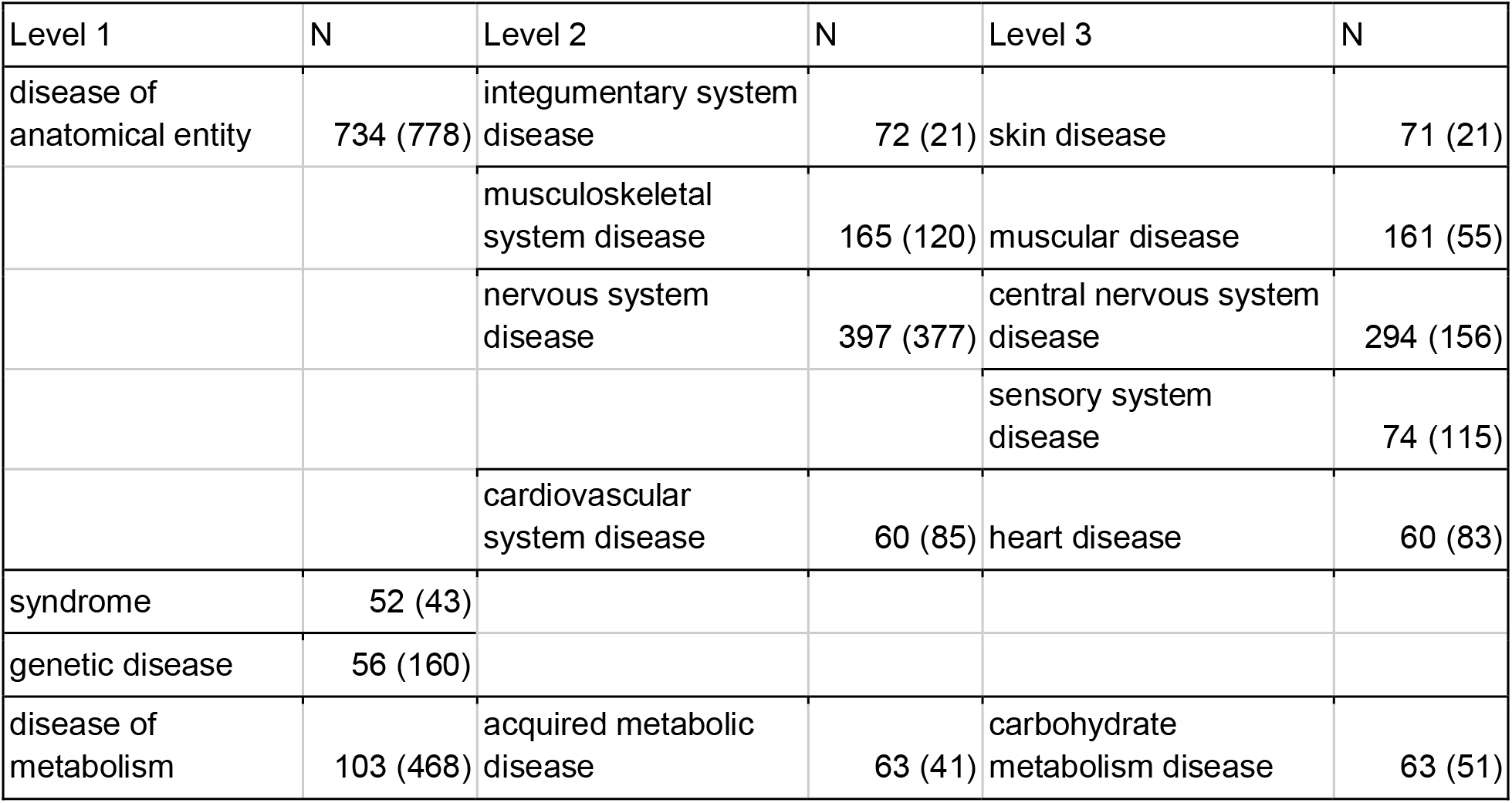
DiseaseOntology term analysis. Only terms responsible for at least 5% of all annotated diseases are shown in the top 3 levels of the ontology. N is the number of mutations disrupting coiled-coil structures (in parenthesis expected values are shown).

| Level 1 | N | Level 2 | N | Level 3 | N |
| --- | --- | --- | --- | --- | --- |
| disease of anatomical entity | 734 (778) | integumentary system disease | 72 (21) | skin disease | 71 (21) |
|  |  | musculoskeletal system disease | 165 (120) | muscular disease | 161 (55) |
|  |  | nervous system disease | 397 (377) | central nervous system disease | 294 (156) |
|  |  |  |  | sensory system disease | 74 (115) |
|  |  | cardiovascular system disease | 60 (85) | heart disease | 60 (83) |
| syndrome | 52 (43) |  |  |  |  |
| genetic disease | 56 (160) |  |  |  |  |
| disease of metabolism | 103 (468) | acquired metabolic disease | 63 (41) | carbohydrate metabolism disease | 63 (51) |

## Discussion

Several studies investigated the structural consequences of inherited disease-causing mutations^16^, one of them also including coiled-coils^14^. In this manuscript, we used four different predictors to assess the structural consequences of variations in coiled-coils, then extended our analysis by incorporating structural data and features responsible for the proper assembly of coiled-coil complexes. We found that DMs accumulate in heptad positions critical for the assembly of coiled-coils (in line with the findings published by Mohanasundaram *et al.^14^*), N-terminal parts of coiled-coils are more abundant in DMs, and mutations mostly strongy affect homooligomeric coiled-coils.

Sequence properties are often used to characterize substitutions, as they often can be connected to structural changes. In this case, grouping amino acids based on their possible role in coiled-coil formation highlights the critical role of charged residues. While mutations on hydrophobic residues impair coiled-coil structures to the same extent as in the case of globular proteins, charge changes often perturb coiled-coil formation. Notably, not only the change of net charge influences coiled-coils, but residues bearing negative charges also do not seem to be interchangeable. This effect is attributable to the helix formation tendency of glutamic acid^17^ that was also proposed in the case of single-α helices^18^. The most characteristic feature of coiled-coils is their repeated register position pattern. Steric clashes and loss of hydrophobic interactions dominate in ‘a’ and ‘d’ positions, while the loss of electrostatic interactions mostly occurs in ‘e’ and ‘g’ positions. Outward-facing residues can also carry essential roles sometimes: they can serve as outside staples that stabilize the alpha-helix by electrostatic interactions, or they can provide a binding site for other molecules (Figure 6, top). For example, the ubiquitin-binding domain (UBAN), conserved in optineurin (OPTN) is part of a coiled-coil, specifically recognizing ubiquitin chains binding to the accessible surface of the coiled-coil^19^. The nuclear factor-κB (NF-κB) pathway plays an important role in regulating inflammation, adaptive and innate immune responses, and cell death via transcriptional targets, such as IL-1β^20^. In the canonical pathway, NF-κB factors are retained in an inactive state via binding to OPTN^19^. The E478G mutation in the UBAN of OPTN abolishes its NF-κB suppressive activity^21^, as residues involved in linear ubiquitin-binding correspond to the residues crucial for keeping NF-κB inactive^22^. The mutations result in significant up-regulation of IL-1β, causing neuroinflammation and neuronal cell death of motor neurons, leading to Amyotrophic Lateral Sclerosis^23^.

**Figure 6:**
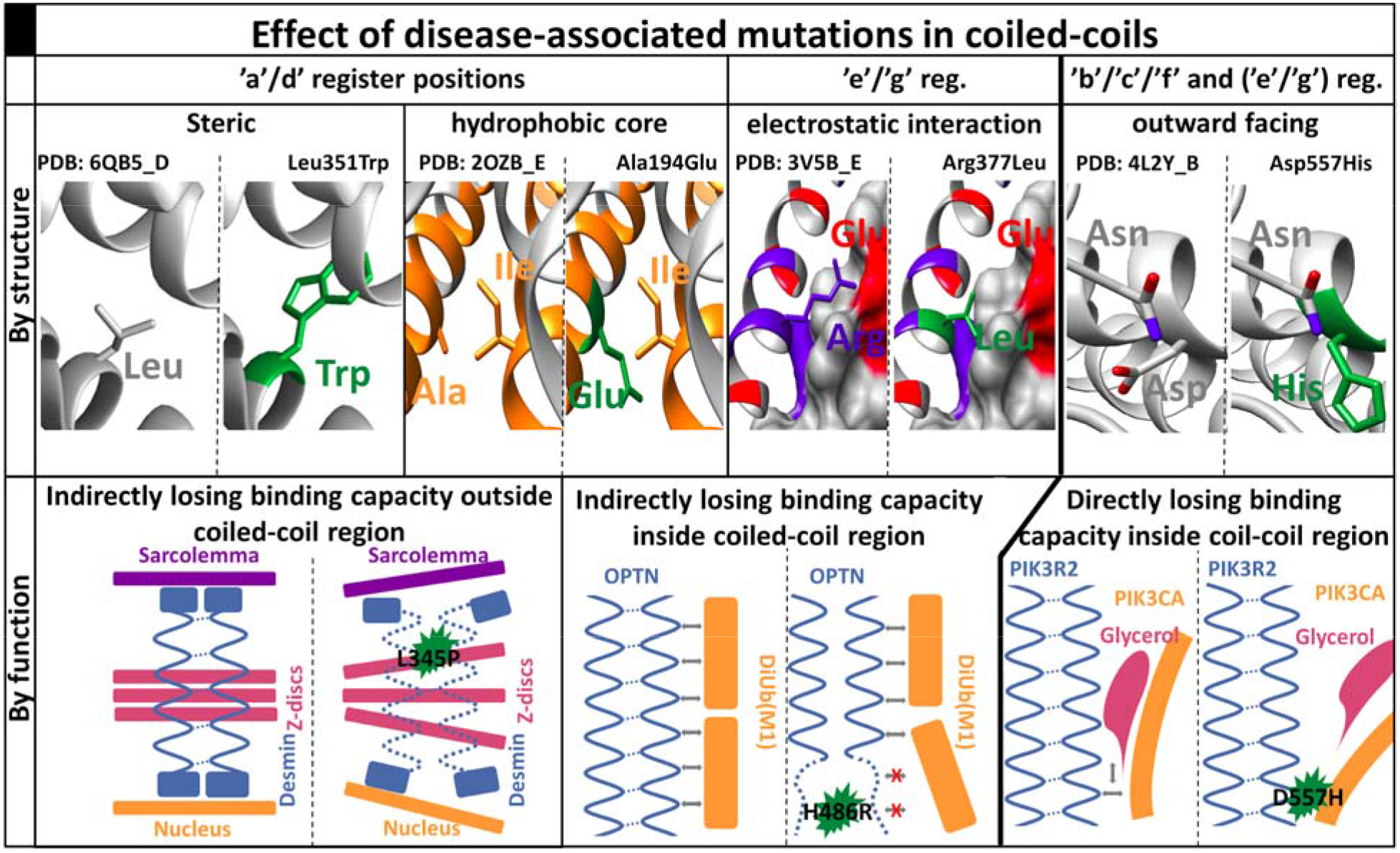
Flavors of disease-causing mutations in coiled-coils. Top row: the structural effect of variations (the ‘Rotamers’ function of Chimera were used to visualize the mutation). Bottom row: functional consequences of mutations. Left side: structure/function in healthy people; Right side: perturbed structures/function having pathological condition. Green color indicates disease-causing mutation; On the structure, orange: hydrophobic; blue: positive; red: negative.

From a functional point of view, mutations falling into distinct structural categories may have different effects. DMs harboring residues contributing to the hydrophobic core usually have an indirect consequence (Figure 6, bottom). In the first scenario, the effect of the mutation manifests outside the coiled-coil region. Desmins are large scaffolding proteins connecting the Sarcolemma, Z-discs, and the nucleus^24^. They consist of elongated coiled-coil regions, with a head and tail unit at their termini. Mutations in the coiled-coil regions disrupt the coiled-coil structure (e.g, DESM: L345P), eventually leading to the disorganization of Z discs and affecting the integrity of the cellular IF network^25^. Mutations often impair coiled-coils directly, so they lose (some of) their binding affinity to molecules interacting with them. The H486R mutation in OPTN perturbs the structure of the UBAN domain and causes low-grade inflammation that leads to glaucoma^26^. However, in contrast to other mutants that were shown to have a direct role in interacting with ubiquitin, this residue points inside the coiled-coil, and only reduces the binding affinity^27^. In the third scenario, mutations are occurring in the coiled-coil, however on a residue facing outward. An example of disruption of direct binding is the mutation affecting interaction of PIK3CA-PIK3R2-glycerol complex. PIK3R2 possesses a two-stranded coiled-coil and forms a heterodimer regulatory unit with PIK3CA via H-bond between N345 of PIK3CA and D557 of PIK3R2. Their complex structure also preserves a groove, providing a room for binding glycerol^28^, which is perturbed by the D557H mutation. The lost direct contact of Asp sidechain with the glycerol, as well as the lack of negative charge positioning the molecule (which is abolished with the positively charged and larger histidine) were proposed to impact binding negatively^29^. PIK3R2 was associated with Megalencephaly-Polymicrogyria-Polydactyly-Hydrocephalus^30^, although the molecular details of the disease were not revealed yet. Besides perturbing structural stability and folding leading to toxic conformations, mutations may also modulate degradation or lead to improper trafficking ^31^. For example, assembly of the Non-POU domain-containing octamer-binding protein is mediated via antiparallel coiled-coil domains and single-α helices^32,33^. The R293H mutation in the coiled-coil domain was shown to lead to subnuclear mislocalization and resulting in endocrine-related tumors^34^.

The different types of coiled-coils utilizing different strategies to achieve a folded state. Many studies already suggested that highly conserved sequence patterns (so-called trigger sequences) are responsible for initiating coiled-coil assembly: for example a seven residue highly conserved motif is required for the folding of the Human Macrophage Scavenger Receptor oligomerization domain^35^, and germline mutation in this region is associated with prostate cancer risk^36^. Another way to initialize assembly occurs during co-translation, as in the case of Peripherin including two-stranded parallel coiled-coils^37^, which also accomodate a disease-causing mutation at the N-terminal region in one of it’s coiled-coil regions^38^. A possible interpretation for the N-terminal accumulation of DMs might lie in the co-translational initiation of the folding and stabilization of α-helices as they emerge from the ribosome^39^. Although abolishing the process likely affects superhelix assembly, this phenomenon only serves as an explanation for mutations in parallel coiled-coils. The critical role of terminal regions are also well-marked in antiparallel coiled-coils: SMC1 forms a complex with SMC3 via their globular N- and C-terminal domains. In both proteins the head and tail regions are connected by antiparallel coiled-coils, and most of the identified DMs gather at their beginning/end of the coiled-coil domains^40^. The proposed antiparallel intramolecular coiled-coil of KIF21A gathers several DMs, predominantly occupying the termini of the coiled-coil^41^, responsible for congenital fibrosis. Thus, although the exact molecular background was not revealed yet, there is a number of evidence supporting the critical role of certain segments in coiled-coils (trigger sites or terminal regions), with an underlied role of N-terminal residues.

Oligomer coiled coils are formed via mutually synergistic folding (MSF), which means interacting partners are disordered in monomer form and the folding and the binding process cannot be separated^42^. The energetic spectrum of such complexes is heterogeneous, including strongly bound structures and weak associations too. Coiled-coil proteins form a particular subclass having relatively weak stabilizing energies, where the interaction plays an essential role in forming a stable complex^43^. It is expected that these proteins are often bound by transient interactions. This is somewhat supported by the fact that both neutral and harmful mutations cause a lower average energetic change compared to other proteins of the proteome. Notably, some of these changes may have a double effect not represented in our calculations, as a significant proportion of mutations affect homooligomers, where both subunits may be exposed to the same mutation.

Our results suggest that mutations in coiled-coils are responsible for several muscular, skin, and neurodegenerative diseases. The role of postsynaptic density (PSD) proteins in neural degeneration is often revisited in various papers^44,45^. In our recent article, we showed that postsynaptic density proteins are exceptionally modular, and they are highly enriched in coiled-coil regions^46^. These results further underline the critical role of coiled-coil containing proteins in the PSD, and advocates the importance of modular protein structures in disease development. A handful of popular methods are available to predict the effect of variations^47,48^ or to highlight vulnerable regions in proteins^49,50^, yet most of these are based on purely statistical approaches. Methods incorporating structural information are largely limited to general features of PDB structures, or prediction of transmembrane domains or disordered segments, although no currently available methods incorporate features of coiled-coils. We showed that basic properties of coiled-coils, such as register position, oligomerization state and position along the region significantly influence the formation of coiled-coils. Since coiled-coil region prediction typically has short run times, we suggest that including such data into state-of-the-art predictors to increase their accuracy would be feasible.

## Methods

### Datasets

The human proteome was downloaded from UniProt^51^, germline variations were obtained from humsavar^51^ (Supplementary Table 1). For redundancy filtering CD-HIT^52^ was applied on the human proteome in an incremental manner, filtering identical proteins to 90, 70, 50 and finally to 40% identity using 5,4,3 and 2 word lengths, respectively (Supplementary Table 2).

### Coiled-coil predictions

Coiled-coil regions were determined using DeepCoil^53^, MarCoil^54^, Ncoils^55^ and Paircoil^56^ (Supplementary Table 3), applying default cutoff values suggested in their descriptive articles. In the case of DeepCoil we utilized the ‘PSSM’ flavor: we generated PSSM for each sequence, using PSI-BLAST with three iterations and 10^−5^ e-value cutoff on the SwissProt database. Coiled-coil positions were predicted using MarCoil, Ncoils and Paircoil (Supplementary Table 4). Oligomerization states were defined using LogiCoil (Supplementary Table 4). Single-α Helix regions^57^ were used as a filter, to reduce false positive hits (Supplementary Table 5).

### Statistical tests

χ^2^ tests were performed in contingency tables (Table 2). Odds ratios were defined as:

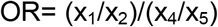

Enrichments on Supplementary figure 1 were defined as:

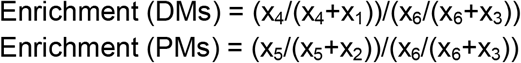

χ^2^ was applied to find the significance of the relation between DMs and coiled-coils (Supplementary Table 6) and the significant importance of the first seven residues of coiled-coil sequences (Supplementary Table 7).

Kolmogorov-Smirnov tests were used to estimate the significance of the distribution of mutations at the first 28 residues of coiled-coils (Supplementary Table 8) and along the coiled-coil sequences (Supplementary Table 9).

χ^2^ test was performed to find the significance of distribution of DMs into coiled-coils with different lengths (Supplementary Table 10)

To estimate the significance of residue changes, we eliminated the sporadic error of the data by performing bootstrap analysis. We randomly selected 80% of the data 100 times and the significance was determined by calculating the average and standard deviations of the data according to the 68-95-99.7 rule (Supplementary Table 11-12).

χ^2^ was used to find the significance of the distribution of DMs into different coiled-coils positions (Supplementary Table 13)

All tests and analysis were performed to the different predictors separately. To produce figures, in each case, we calculated the mean of different predictors (Supplementary Table 6-13). Although we plotted mean values, if predictors showed different trends, we noted it in the text.

### DiseaseOntology term analysis

Disease ontology terms were mapped using MIM identifiers from humsavar and DiseaseOntology^58^. Only identifiers linked to DMs, where all methods predicted coiled-coil were used. For the analysis the top three level of the ontology was applied, and the number of mutations were counted in each disease category - only terms occurring in coiled-coil containing proteins are shown in Supplementary Table 14, and only terms responsible for at least 5% of all annotated diseases shown in Table 1. Next we mapped all mutations in a similar manner. Expected values were calculated by normalizing these numbers on each term with the proportion of all coiled-coil mutations.

### Assigning structures to amino acid sequences

We used BLAST on sequences from the non-redundant human proteome against the PDB with 10^−5^ e-value. Chimeric proteins were discarded. We used the greedy algorithm to select structures with 100% identity, with the most variations mapped on them (Supplementary Table 15). On all PDB structures we considered biomatrix transformations as defined in the PDB files to detect all possible coiled-coils.

### Calculating structural and energetic properties

We detected coiled-coils using SOCKET^59^ with default settings. Coiled-coil features (heptad positions, the number of strands, angle of strands (Supplementary Table 15-16)) were determined based on SOCKET output. For monomer/homooligomer/heterooligomer assignment, we checked which BLAST query corresponds to the detected coiled-coil regions (Supplementary Table 18). Energy calculations were performed using FoldX^60^. ΔΔ*G* calculations were executed on previously optimized structures and were performed five times. All reported ΔΔ*G* values represent the average of these independent runs. In 76 cases (less than 1%) we experienced problems with FoldX, these cases were omitted (Supplementary Table 19). Calculated structural features shown on Figure 4 are based on values from Supplementary Table 20. Energetic changes on figure are categorized as highly stabilizing (<−1.84□kcal/mol), stabilizing (− 1.84 to − 0.92□kcal/mol), slightly stabilizing (− 0.92 to − 0.46□kcal/mol), neutral (−0.46 to + 0.46□kcal/mol), slightly destabilizing (+ 0.46 to + 0.92□kcal/mol), destabilizing (+ 0.92 to + 1.84 kcal/mol) and highly destabilizing (> + 1.84 kcal/mol).

### Visualization

Images were prepared using UCSF Chimera^61^.

