## Supplementary Material for "Distribution of disease-causing germline mutations in coiled-coils suggests essential role of their N-terminal region"

### Supplementary Figures

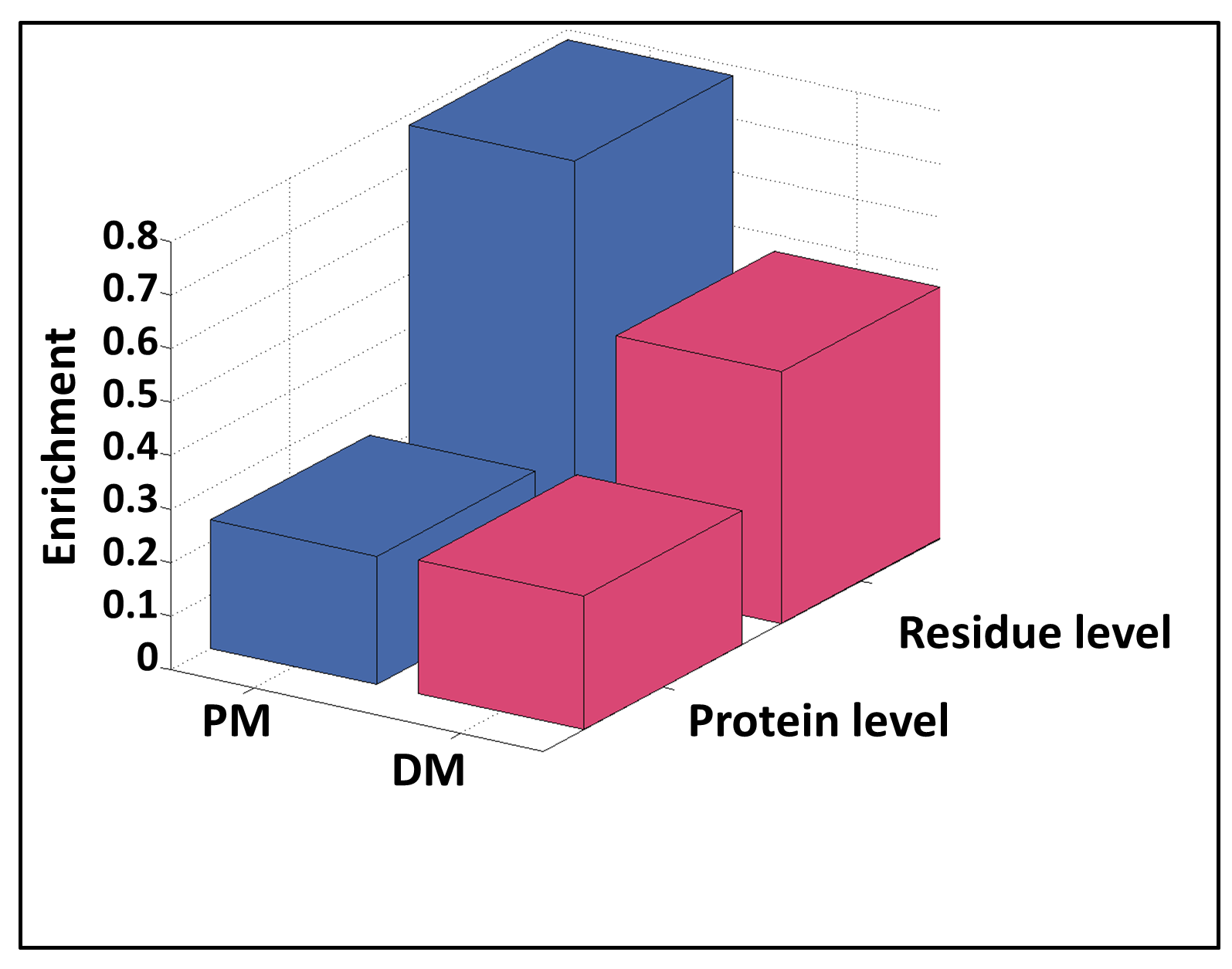

***SFigure 1: Relation between DMs and coiled-coils.*** *A) Enrichment of PMs (blue) and DMs (red) in proteins and in residues.*

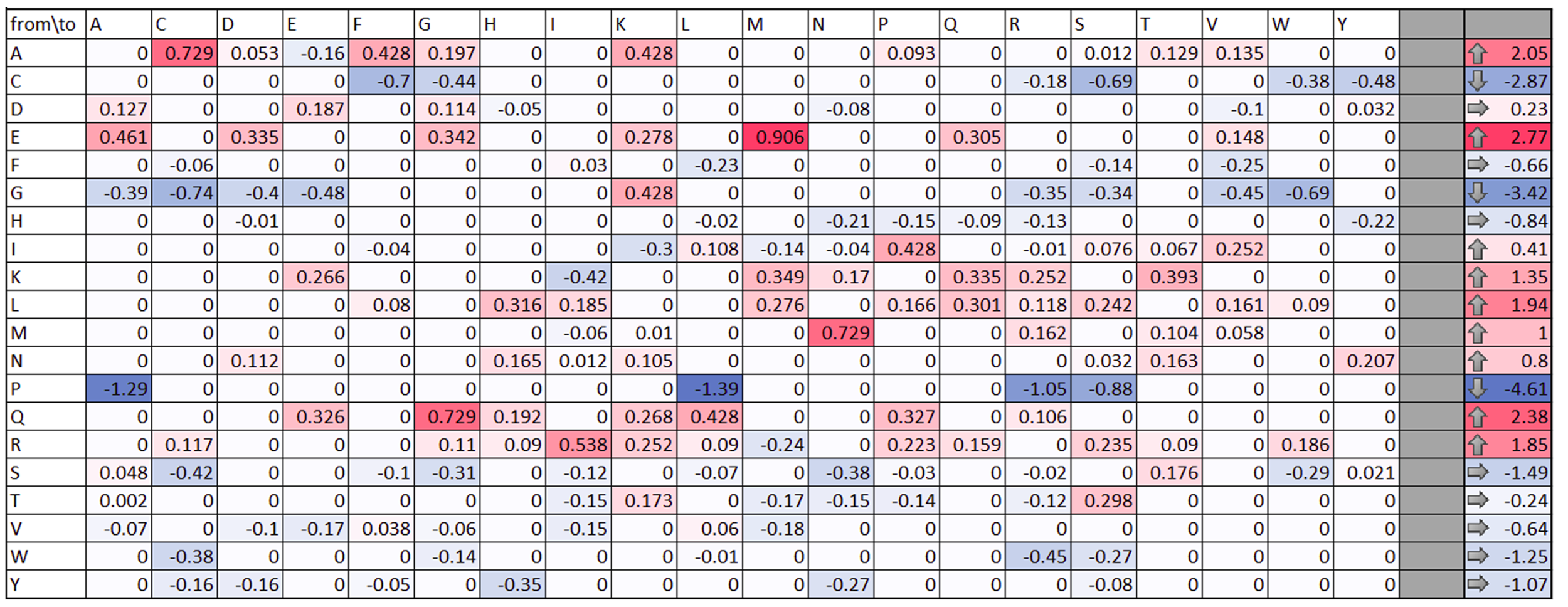

***SFigure 2: Amino acid changes in coiled-coils.*** *Left) Residue change preferences by DMs in the proteome (negative values, also marked with the shades of blue) and in coiled-coil regions (positive values, also marked with the shades of red). Values show the logarithm of ratio of DMs changing given residues types Right) Targeted residue type preferences by DMs in the proteome (negative values) and in coiled-coil regions (positive values).*

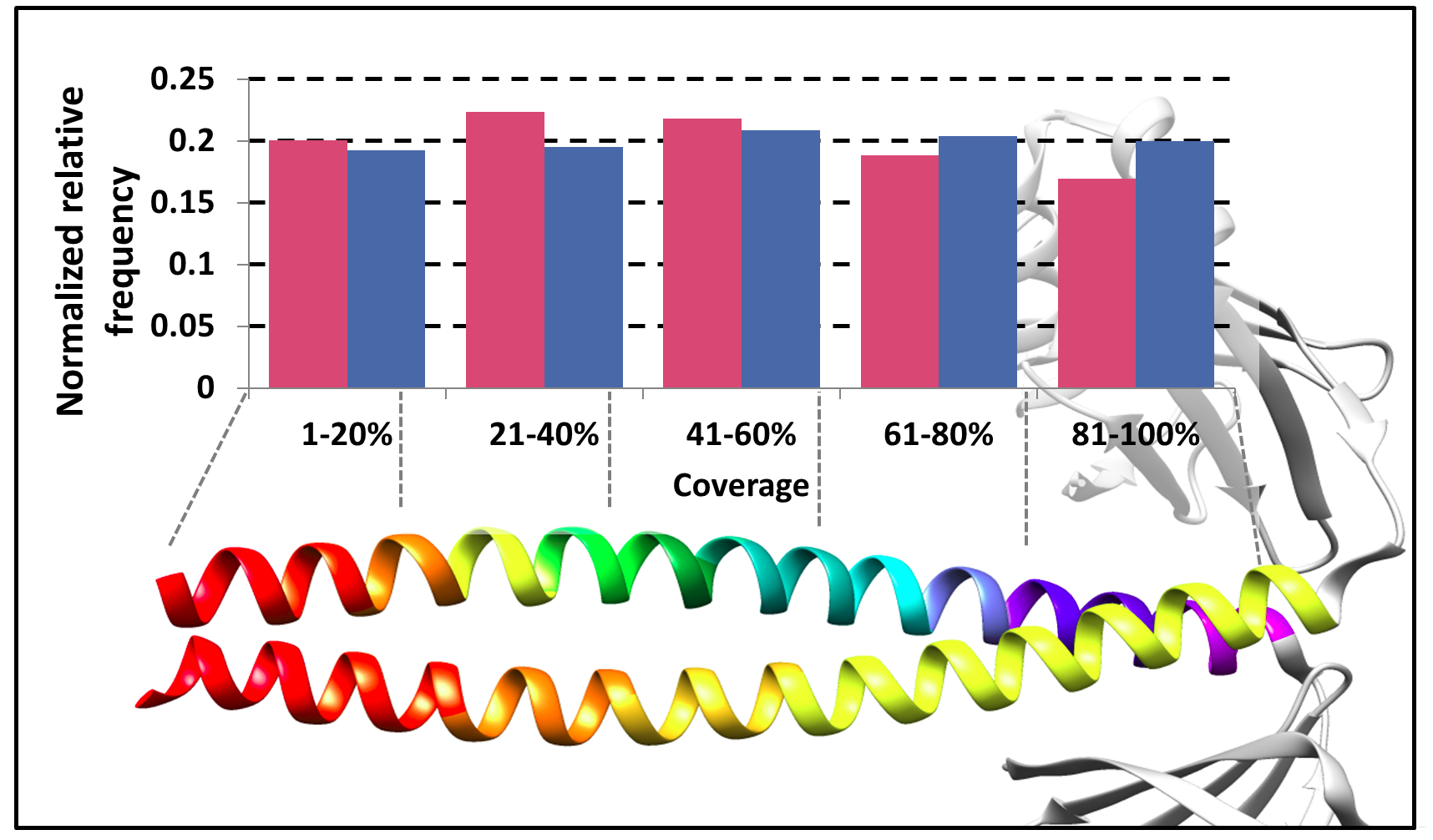

***SFigure 3: Variations along coiled-coil segments.*** *Distribution of variations in the sequence. X-axis shows the coverage of the sequences.*
